# An immunoresponsive three-dimensional urine-tolerant human urothelial (3D-UHU) model to study urinary tract infection

**DOI:** 10.1101/2022.07.22.501108

**Authors:** Nazila V. Jafari, Jennifer L. Rohn

**Affiliations:** Division of Medicine, University College London, Royal Free Hospital Campus, London, UK

**Keywords:** Bladder 3D model, CD markers, uroplakins, cytokeratins, tight junctions, innate immunity

## Abstract

Murine models of urinary tract infection (UTI) have significantly improved our understanding of host-pathogen interactions. However, given some differences between the rodent and human bladder which may modulate bacterial response, including certain biomarkers, urothelial thickness and the concentration of urine, the development of new human-based models is important to complement mouse studies and to provide a more complete picture of UTI in patients. We originally developed a human urothelial three-dimensional (3D) model which was urine tolerant and demonstrated several biomarkers of the urothelium, but it only achieved human thickness in heterogenous, multi-layered zones and did not demonstrate the comprehensive differentiation status needed to achieve barrier function. Here, we report an improved 3D urine-tolerant human urothelial (3D-UHU) model, which after 18-20 days of growth, stratifies uniformly to 7 layers. We confirmed by flow-cytometric analysis that the model is comprised of the three expected, distinct human cell types. The apical surface differentiated into large, CD227^+^ umbrella-like cells expressing uroplakin-1A, II, III, and cytokeratin 20, all of which are important terminal differentiation markers, and a glycosaminoglycan layer. Below this layer, several layers of intermediate cells were present, with a single underlying layer of CD271^+^ basal cells. The apical surface also expressed E-cadherin, ZO-1, claudin-1 and -3, necessary for barrier formation; barrier integrity was confirmed by transepithelial electrical resistance and FITC-dextran permeability assay. Infection with both Gram-negative and Gram-positive bacterial classes elicited elevated levels of pro-inflammatory cytokines and chemokines characteristic of urinary tract infection in humans, and caused a decrease in barrier function, suggesting that the 3D-UHU model holds promise for studying host-pathogen interactions and host innate immune response.

## Introduction

The urinary tract is the most common site of bacterial infection in humans, and urinary tract infection (UTI) affects ∼150 million people around the world each year. Approximately 40% of women and 12% of men experience a symptomatic UTI during their lifetime, although infants and children are also susceptible. While acute UTIs are usually self-limiting, within 6-12 months, a quarter of affected women will suffer recurrent UTI despite treatment^1,2^.

The most apparent function of the urinary bladder is to store and void large volumes of urine. It is comprised of three tissue layers: the luminal urothelium, the lamina propria, and the detrusor muscle. The urothelium is a stratified, transitional epithelium that lines the surface of the renal pelvis, ureters, urinary bladder and proximal urethra. It serves as a barrier that prevents the diffusion of toxic substances such as acid and urea and defends against pathogens from the external environment^3,4,5^. The urothelium consists of three cell types: the undifferentiated basal cells, intermediate cells and fully differentiated luminal umbrella cells^6^. Forming a single layer along the basement membrane, the basal cells are the smallest (5-10 µm in diameter) and are the most abundant cell population in adult urothelium^4,7^. The intermediate cells lie directly above the basal cells and are considerably larger (20 µm in diameter). In the urinary bladder of humans there are up to five intermediate cell layers, whereas in rodents only one layer is present^3,7,8^. Facing the luminal surface are large (50-120 µm in diameter), hexagonal and highly specialized cells known as superficial or umbrella cells^4,9^.

The apical surface of the umbrella cells is covered by a crystalline lattice comprising four key uroplakin proteins (UPK1A, UPK1B, UPKII, UPKIII) that together form a unique asymmetric unit membrane (AUM) plaque^4^. Uroplakins are considered to be terminal differentiation markers of the urothelium. They initially form heterodimers (UPK1A-UPKII and UPK1B-UPKIII), then the resulting moieties interact to form heterotetradimers. Six heterotetradimers assemble a 16 nm protein particle arranged into a hexagonal lattice and are presented to the apical membrane which decorates up to 90% of the luminal surface^9^. In rodents, uroplakins are expressed within all urothelial layers, whilst in humans they are primarily expressed in the umbrella cells, highlighting species specificity in their expression pattens^10^.

The urothelium also expresses several cytokeratins including cytokeratin 5 (CK5), CK7, CK8, CK13, CK14, and CK20. Cytokeratin expression varies in the urothelium depending on its location and some are expressed in a differentiation-specific manner, such as CK20 which is highly expressed in the umbrella cells^7,8^. Despite the multilayer structure of the urothelium, the outermost umbrella cell layer forms the barrier to pathogens, urine and its associated metabolites, solutes, and water. This barrier itself is multifaceted, comprised of the apical membrane, the umbrella cell tight junction (containing proteins such as occludins and claudins), and the glycosaminoglycan (GAG) layer covering the umbrella cells^11,12^.

The impermeable barrier of the umbrella cells together with protective glycan layer and frequent unidirectional flow of urine discourages bacterial adherence to the urothelium. Other factors such as changes in urine osmolarity, pH, soluble IgA, uromodulin (Tamm-Horsfall urinary glycoprotein), iron chelating siderophores and antimicrobial peptides (AMPs) can additionally limit bacterial attachment to urothelia^13^. Furthermore, the urothelium employs another defence mechanism to reduce bacterial load by undergoing cell death and cell exfoliation into the urine. This allows the removal of cells that are associated with adherent and intracellular bacteria^14,15^.

The urothelium also expresses multiple toll-like receptors (TLRs) which recognise pathogen-associated molecular patterns (PAMPs)^16,17,18^. The activation of TLRs triggers the production of inflammatory mediators such as cytokine and chemokines. The common TLRs identified in the urinary tract include TLR-2, TLR-3, TLR-4, TLR-5, and TLR-9, (TLR-11 only in mice)^19^. Studies have reported that normal human urothelium express TLR-5 weakly, TLR-2, TLR-3, and TLR-7 moderately, and TLR-4 and TLR-9 strongly^20,21^.

Although murine models of UTI have proven to be a powerful tool for advancing our understanding of UTI pathogenesis^22^, mouse models have limitations. Physiological, structural, and genetic differences can limit how well mice reproduce key aspects of disease pathogenesis or pathogen’s ability to replicate causing human-like diseases^23^. Development of new treatments not only requires an in-depth understanding of disease pathogenesis but importantly, access to appropriate model systems to study host-pathogen interactions. While *in vitro* cell models are limited in terms of immune response and systemic crosstalk, their human microenvironmental context nevertheless can provide complementary information to animal studies^24^. Indeed, biomimetic *in vitro* models of the human urothelium have advanced significantly in recent years, offering vital information, but most are short-lived when in a urine environment^25,26^. This is a significant disadvantage, as urine is the natural context in which uropathogens interact at the host cell interface. We overcame this problem in our previous model^27^, but the urothelium was non-homogenous in terms of differentiated surface coverage, and possibly as a result, the barrier function was not robust. Here, we report an improved fully urine-tolerant human urothelium that can stratify to a human urothelial thickness of seven layers, exhibits the key biomarkers, possesses good barrier function and secretes appropriate human cytokines in response to infection.

## Results

### An enhanced 3D-UHU model mimics human bladder epithelia

HBLAK cells are a commercially available human bladder epithelial progenitor cells that have been spontaneously immortalized but still retain the ability to terminally differentiate in response to the correct stimuli. Cell cultures grown to a 70-90% confluency were seeded onto a 12-well transwell inserts with CnT-PR medium; when confluent, the cell culture medium was replaced with CnT-PR-3D to initiate differentiation. 15-16 h post-incubation, CnT-PR-3D was replaced with filter-sterilised human urine in the apical compartment. Cultures were maintained with regular media and urine change until the 3D cultures were fully stratified and differentiated on days 18-20. To examine the 3D models microscopically, membranes were fixed then removed from the inserts, and stained for various biomarkers.

Confocal microscopy showed a stratified human bladder 3D model achieving up to 7 cell layers thickness (Fig. 1a). The single optical slices showed densely packed, very small basal cells in the basal layer (Fig. 1b) followed by a few layers (4-5) of slightly larger intermediate cells, and large hexagonal umbrella-like cells forming the top layer (Fig. 1c). The umbrella cell layer was homogenous, showing nearly 100% differentiation without the zones of hypertrophy and undifferentiated monolayers seen in our previous version of this model^27^ (data not shown).

**Figure 1.**
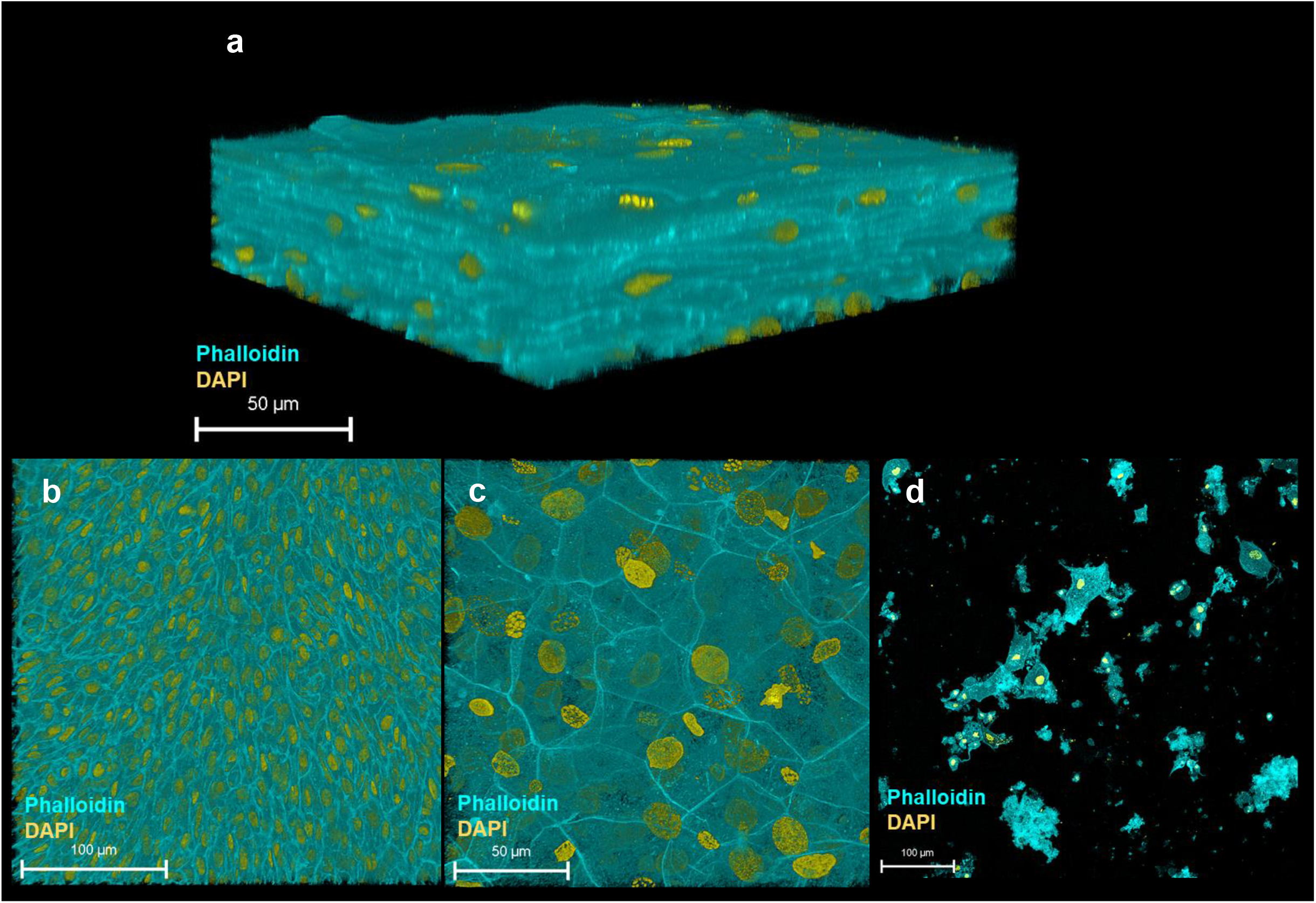
HBLAK cells can differentiate and form a 3D urothelial model, 3D-UHU. (a) 3D confocal image of the urothelial model; (b) single optical slice at the lowest region of the model showing small, tightly packed basal cells; (c) single optical slice at the apical surface exhibiting large, differentiated umbrella-like cells; (d) shed cells detected at day 18 in fully differentiated 3D models (supernatants collected from 12 models). Phalloidin-stained F-actin is presented in cyan and DAPI-stained DNA is presented in yellow. Images are representative of at least three biologically independent experiments.

We examined epithelial cell shedding by collecting spent apical supernatants at regular intervals which were centrifuged onto glass slides prior to fixation and staining. A very low number of shed cells (∼ 25 cells/ well based on nuclei count, rest presumed cell debris) were detected on day 18 onwards with few cells resembling umbrella-like cells (Fig. 1d), suggesting that the model retains most of its umbrella cell layer.

In summary, the 3D-UHU model showed three distinct cell layers with discernible cell size differences and appropriate thickness, reminiscent of a human urothelium.

### The 3D-UHU model expresses human bladder CD cell surface markers

We identified stratified and terminally differentiated 3D-UHU cell types based on their marker expression using flow cytometric analysis. Dissociated single cells were first gated by size and granularity to exclude cellular debris. From the “gated cells”, singlets were sub-gated by their FSC-A/FSC-H properties. Single cells were plotted with the viability dye and cells considered “live” were selected (Fig. 2a-c). From “live” population, cells that were positive for CD9 and CD59 markers were selected (Fig. 2f). CD9 and CD59 have been reported to be expressed by all three cell types: basal, intermediate, and umbrella cells^28^. CD9, CD59 gated cells were plotted with CD44, CD104, and CD271 (Fig. 2d-e). CD4 and CD104 are expressed strongly by basal cells although intermediate cells express these markers in low levels, whereas CD271 is expressed exclusively by basal cells. Intermediate cells have been reported to express a number of CD molecules (e.g., CD9, CD46, CD49b, CD55, CD59, CD95, CD116, CD147)^28^; however, these markers are expressed by basal and umbrella cells to varying degrees of intensity. CD49b and CD95 markers were used to label the intermediate cells (Fig. 2g). To detect the umbrella cells, CD9/CD59 gated cells were plotted with CD47, CD63, and CD227 (Fig. 2h-i). CD47 and CD63 molecules are highly expressed by umbrella cells although they selectively label intermediate and basal cells (Fig. 2h). We identified the umbrella cells by CD227, the marker that is expressed exclusively by these cells (Fig. 2i).

**Figure 2.**
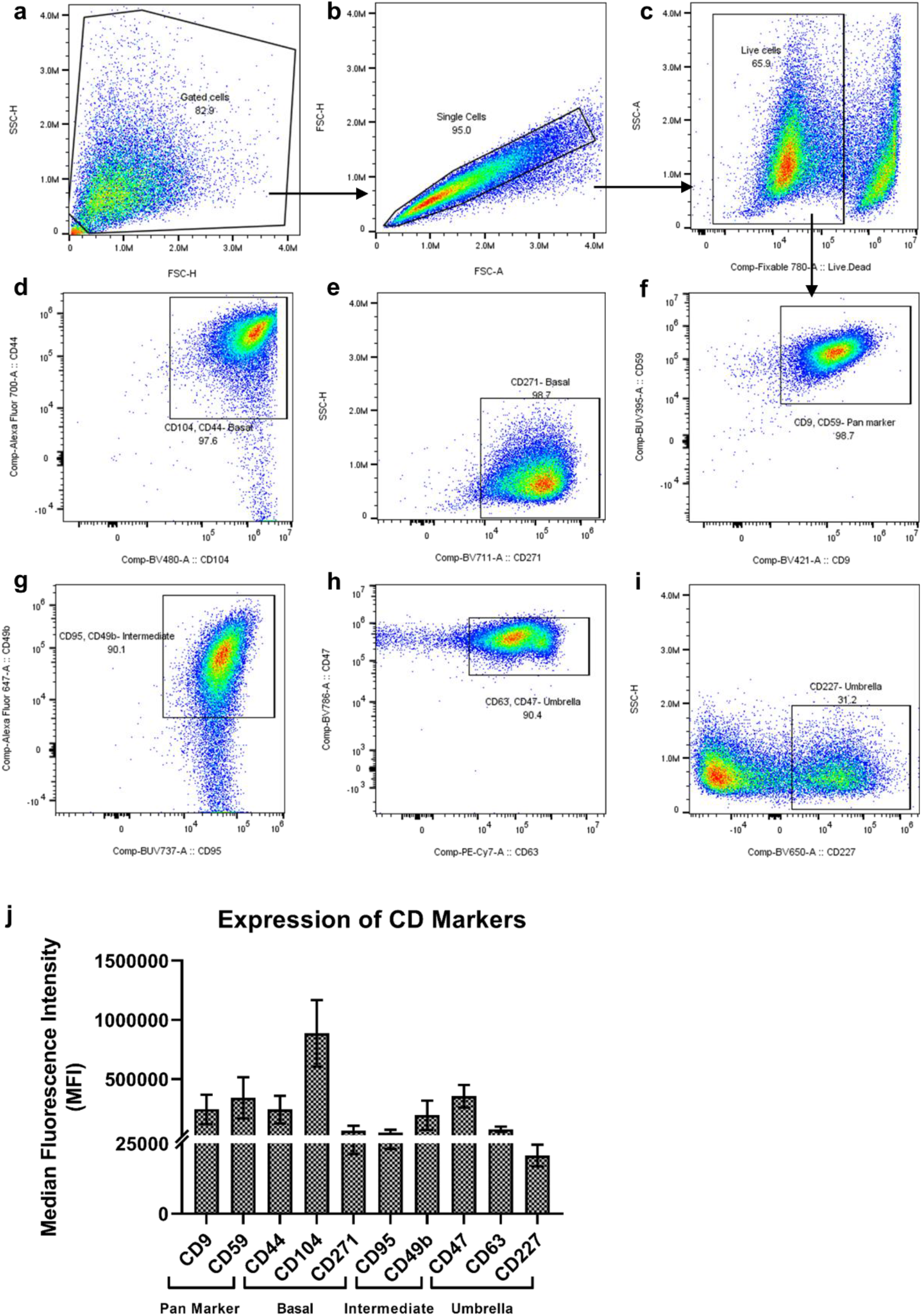
3D-UHU model expresses human bladder urothelial CD phenotypes. (a-c) gating strategy to profile urothelial CD markers by flow cytometry; (d-i) flow cytometry analysis of the expression of CD9/CD59 (pan cell surface markers); CD44/CD104, CD271 (basal cells); CD95/CD49b (intermediate cells); CD63/CD47, CD227 (umbrella cells); (j) expression level (MFI) of CD cell surface markers, data represent mean ± SD, n=4 biologically independent experiments.

We used unstained, fluorescence minus one (FMO), and CD57 marker expressed by urothelial nerve sheath (data not shown) controls to set gate limits and rule out non-specific binding of the panel antibodies to the cellular surface. The median fluorescence intensity (MFI) of CD markers expressed by human bladder 3D model showed a high CD104 and a relatively low CD227 expression (Fig. 2j).

### The 3D-UHU model exhibits human urothelial biomarkers

We characterised the model further by examining the expression of urothelial-specific markers with confocal microscopy. The 3D-UHU model showed the expression of all three uroplakin proteins integral to formation of the AUM. Uroplakin-1A (UPK1A) (Fig. 3a), uroplakin-II (UPKII) (Fig. 3b), and uroplakin-III (UPKIII) (Fig. 3c) were expressed in the umbrella-like cells with UPKIII exhibiting the highest surface expression followed by UPK-1A. Although studies have indicated that cultured urothelial cells do not express UPII, our model showed a modest expression of this marker. Next, we examined the GAG layer expression. The GAG layer consists of glycoproteins and glycolipids and is mainly composed of heparin sulfate, chondroitin sulfate, keratan sulfate, and dermatan sulfate^29^. Although not all of these components were tested due to a lack of commercially available reagents, a chondroitin-rich GAG layer was detected at the umbrella cell layer (Fig. 3d).

**Figure 3.**
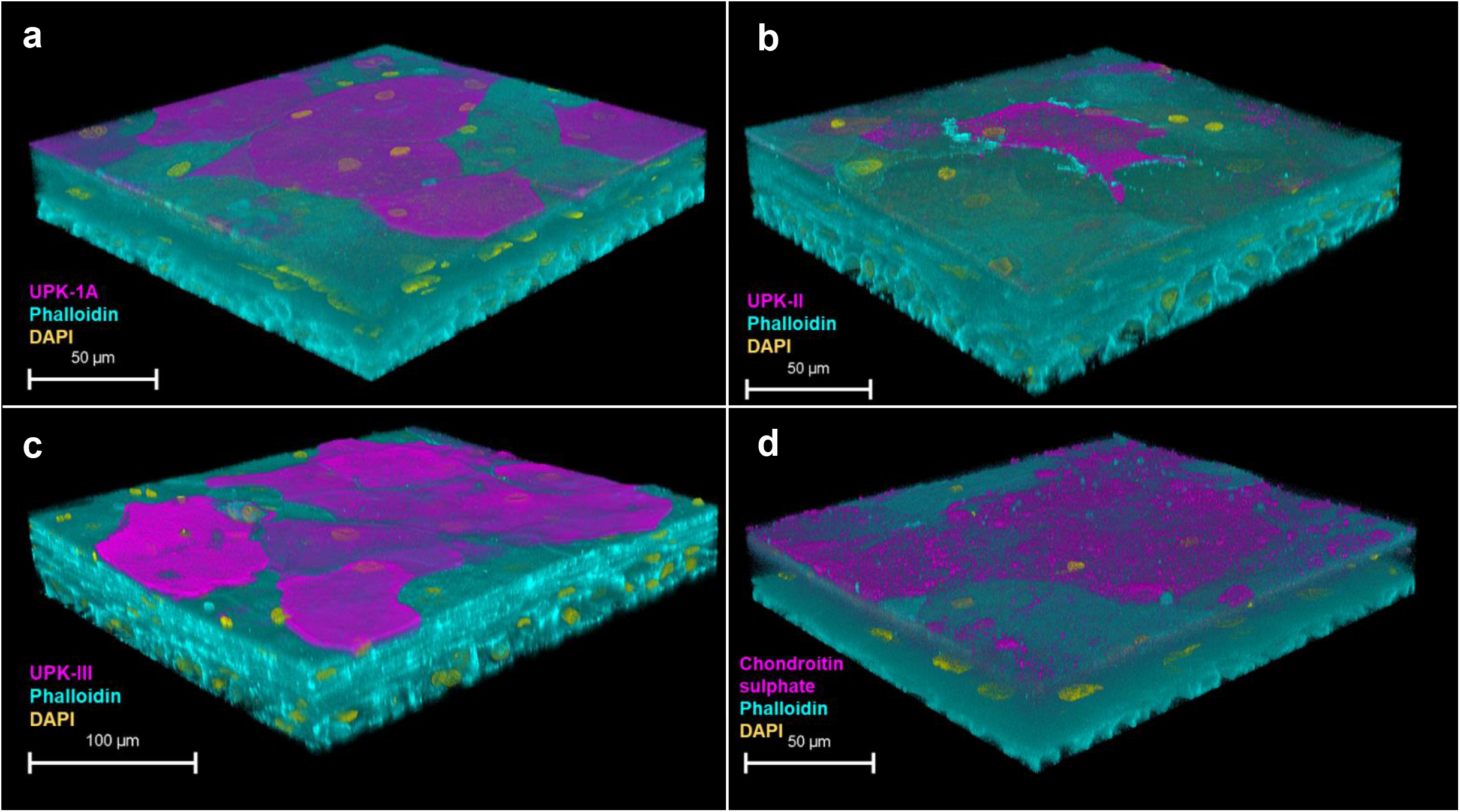
Uroplakin and GAG layer expression of the 3D-UHU model. 3D confocal image of (a) UPK-1A (magenta); (b) UPKII (magenta); (c) UPKIII (magenta); and (d) chondroitin sulphate (magenta) expressed at the apical surface of the 3D-UHU models. Phalloidin-stained F-actin is presented in cyan and DAPI-stained DNA is presented in yellow. Images are representative of at least three biologically independent experiments.

The 3D-UHU also exhibited the correct spatial expression of cytokeratin (CK)8, 13, and 20 (Fig. 4). CK8 was expressed throughout the basal, intermediate, and umbrella cell layers (Fig. 4a). CK13, a late intermediate differentiation marker indicating a switch from basal-like cells to intermediate cells^30,31^ was detected on the late differentiated cell layers (Fig. 4b), while CK20 expression, a terminal differentiation marker was detected only at the apical surface, presented by umbrella-like cells (Fig. 4c).

**Figure 4.**
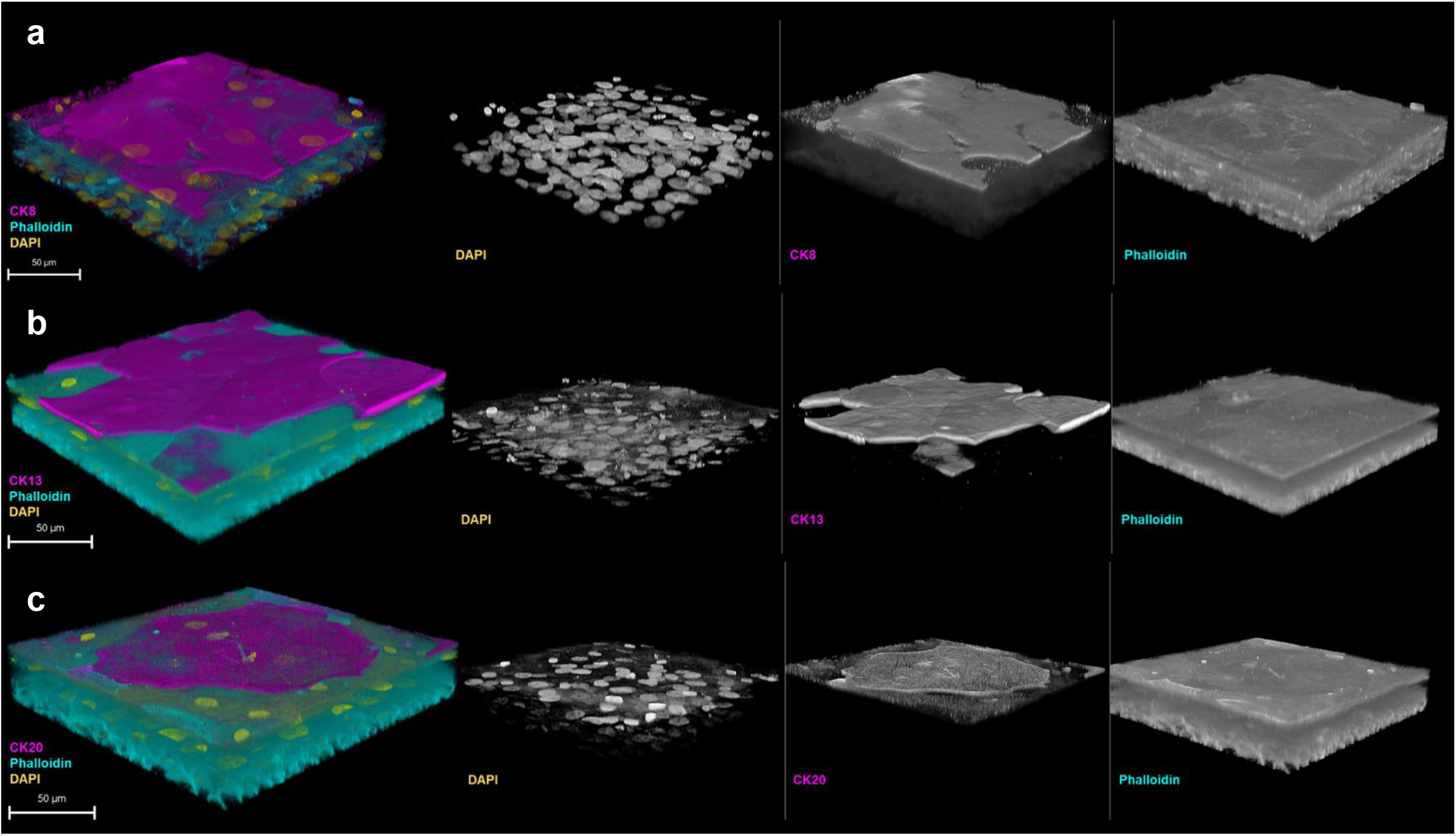
The 3D-UHU model shows cytodifferentiation. 3D confocal image of (a) CK8 (magenta) expressed throughout the layers; (b) CK13 (magenta), and (c) CK20 (magenta) expressed at the late/terminal and terminally differentiated umbrella-like cells, respectively. Phalloidin-stained F-actin is presented in cyan and DAPI-stained DNA is presented in yellow. Images are representative of at least three biologically independent experiments.

### The 3D-UHU expresses adherens and tight junction proteins and forms a barrier

The expression of adherens junction (AJ) E-cadherin and tight junction (TJ) proteins – ZO-1 and claudin 1, 3 – were examined by confocal microscopy (Fig. 5a-f). In the literature, the umbrella cell layer is the only urothelial layer that forms detectable tight and adherens junctions^7^. The 3D-UHU model showed both membranous and cytoplasmic E-cadherin expression in the umbrella-like cells (Fig. 5a). A strong ZO-1 expression was detected in cytoplasm of the umbrella cell layer with a faint staining at cell borders (Fig. 5c), while claudin 1 and claudin 3 exhibited a discontinuous membrane and cytoplasmic staining and the expression was distributed diffusely throughout the strata (Fig. 5e&f).

**Figure 5.**
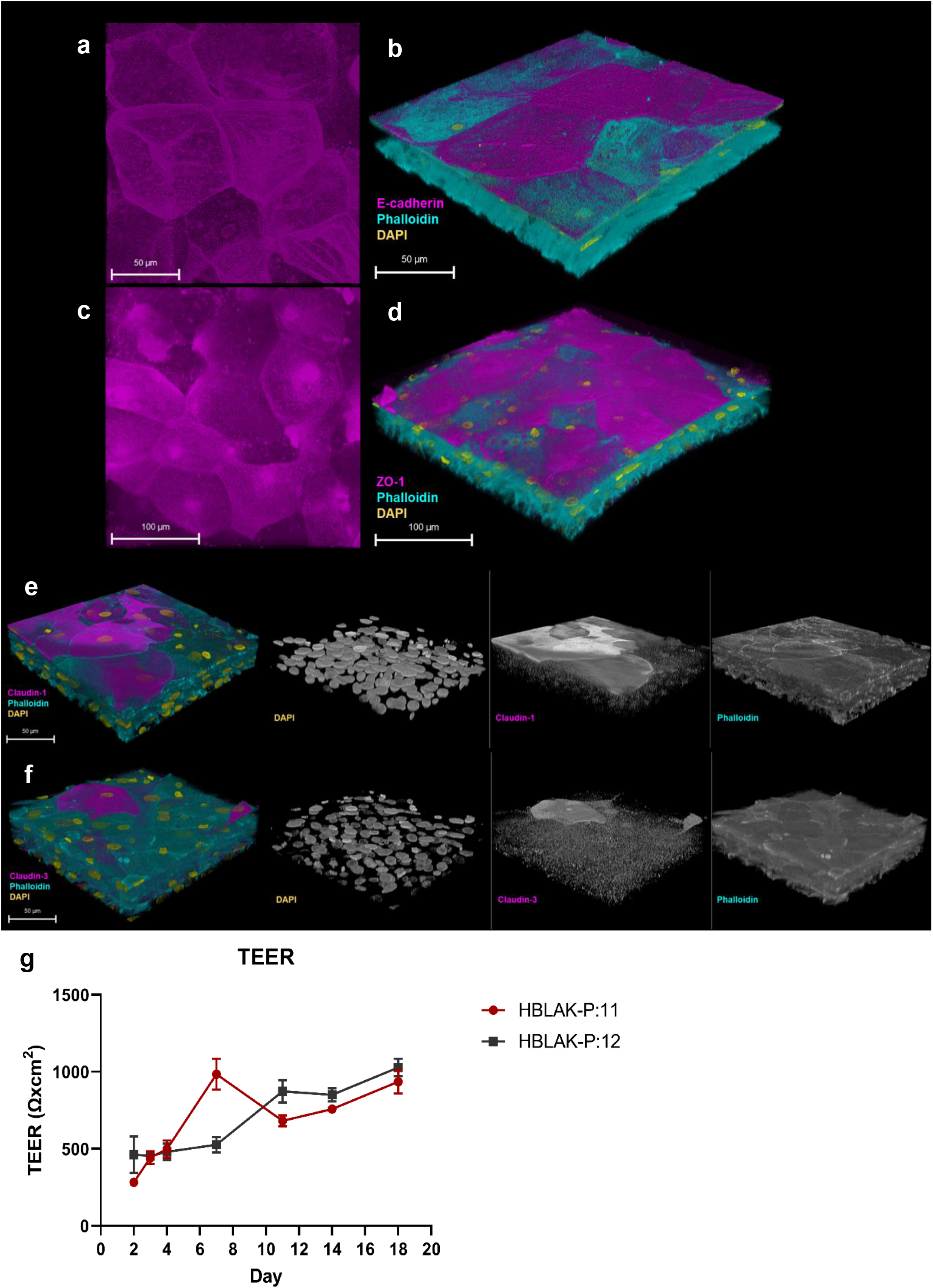
The 3D-UHU model express adherens and tight junction proteins and form a barrier. (a) Single optical slices at apical region of the model showing E-cadherin expression (magenta); (b) 3D confocal image of urothelial model expressing E-cadherin at the apical surface (magenta); (c) Single optical slices at apical region expressing ZO-1 (magenta); (d) 3D confocal image of urothelial model expressing ZO-1 at the apical surface (magenta); (e-f) claudin 1 and claudin 3 presented throughout the layers (magenta). Phalloidin-stained F-actin is presented in cyan and DAPI-stained DNA is presented in yellow. Images are representative of at least three biologically independent experiments. (g) TEER measurement of 3D-UHU model in a time-dependent manner; data represent mean ± SD, n=3 biologically independent experiments.

We determined the barrier integrity by measuring the transepithelial electrical resistance (TEER) in a time-dependent manner (Fig. 5g). TEER was assessed on day 2 (48h post-seeding), day 3 (CnT-PR-3D addition to initiate cell differentiation), and days 4-18 (cell exposure to urine). Although the TEER time-course showed some passage differences around day 8, a stable increase in TEER was detected from day 12 to 18 reaching to ∼1000 Ω.cm^2^.

### HBLAK cells express Toll-like receptors (TLRs)

To examine the expression of TLRs, HBLAK cells were grown to confluency as monolayers on transwell inserts then stained for TLR-2, -4, and -5 (Fig. 6). A moderate to strong expression of TLR-2 and TLR-4 was detected in HBLAK monolayer cells (Fig. 6a&b), while TLR-5 exhibited a low expression (Fig. 6c).

**Figure 6.**
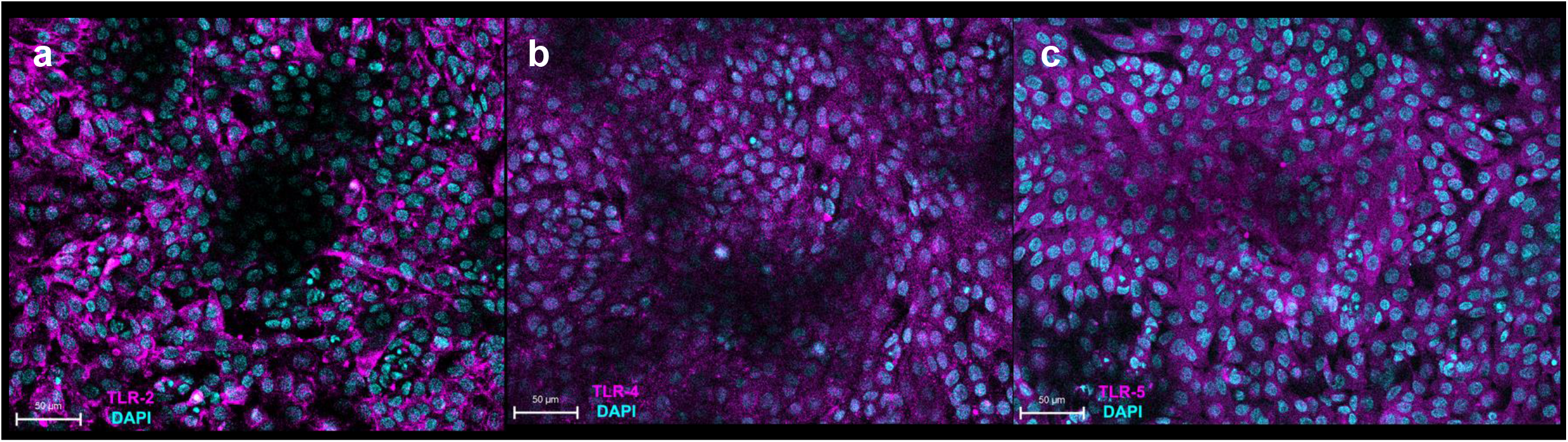
HBLAK cells express toll-like receptors. HBLAK confluent monolayer cells express (a) TLR-2 (magenta); (b) TLR-4 (magenta); (c) TLR-5 (magenta). DAPI-stained DNA is presented in cyan. Images are representative of at least three biologically independent experiments.

### Uropathogenic strains show variation in the disruption of the epithelial barrier

Next, we wanted to evaluate the effect of uropathogens on barrier integrity by assessing the paracellular permeability (Fig. 7). A FITC-dextran (4 kDa) marker was used to measure tight junction permeability in response to uropathogenic *Escherichia coli* CFT073, *E. coli* 83972 (asymptomatic bacteriuria [ABU] isolate), *Enterococcus faecalis* 36 and *E. faecalis* 77 clinical isolates, and *Streptococcus agalactiae* (multiplicity of infection; MOI 10). At 5h post-infection, a similar but slight disruption in barrier integrity was detected compared with uninfected control cells. CFT073 showed a significant (adjusted *p*-value <0.0001) disruption at 24h post-infection compared with uninfected control while *S. agalactiae* had the least effect. *E. faecalis* 36 exhibited a similar effect (adjusted *p*-value 0.0025) on tight junctions as CFT073. *E. coli* 83972, *E. faecalis* 77, and *S. agalactiae* showed a barrier disruption that was not statistically significant compared with uninfected control.

**Figure 7.**
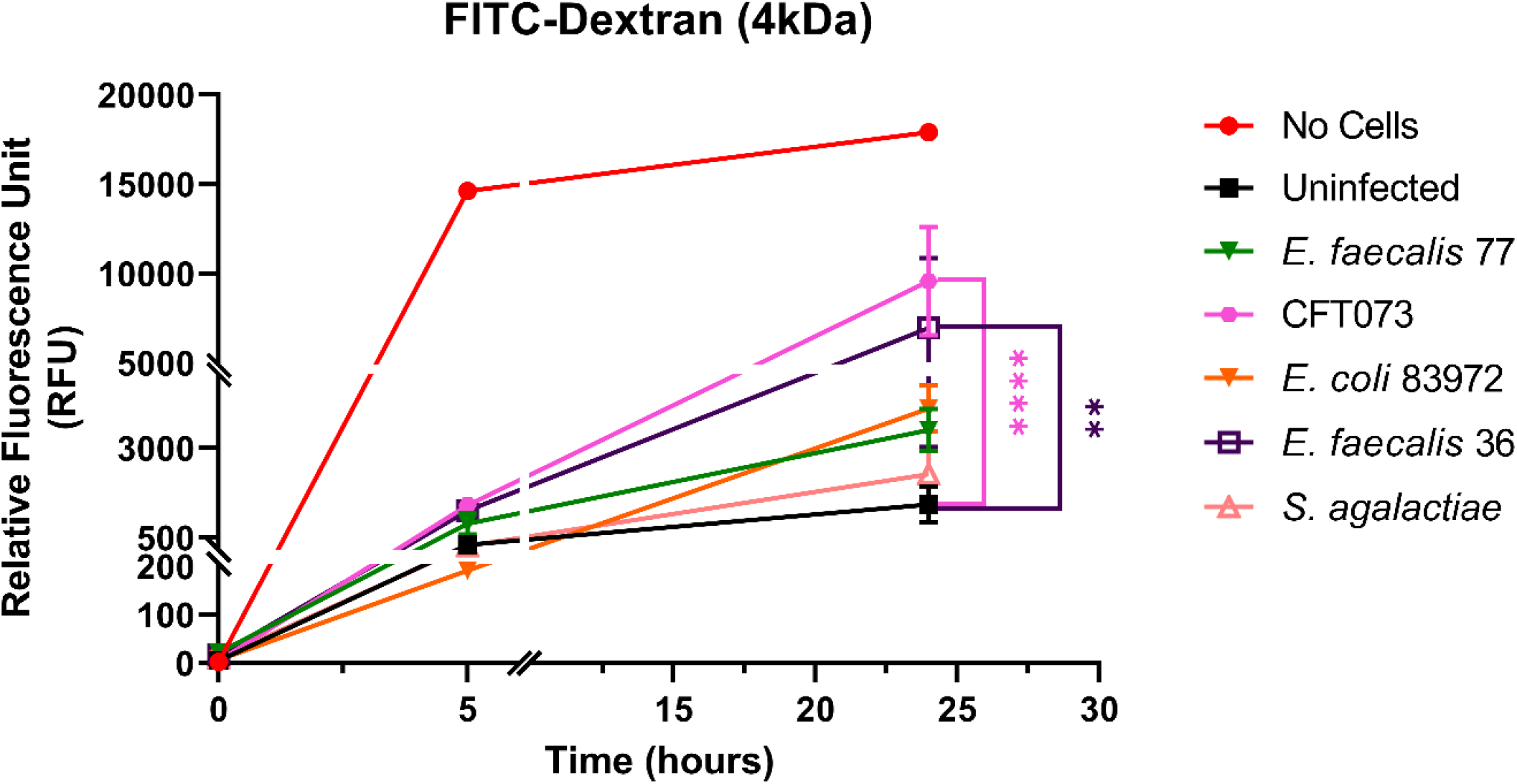
Uropathogens disrupt barrier integrity of 3D-UHU models. The barrier integrity in response to uropathogens (MOI 10) was measured by assessing the FITC-dextran influx at 5h and 24h post-infection. Data represent mean ± SD, n=3 biologically independent experiments. *P* values were obtained using ANOVA with Dunnett’s multiple comparisons test.

### Uropathogenic strains trigger an innate immune response

To determine the 3D-UHU innate immune response to uropathogenic *E. coli* UTI89 and *E. faecalis* 36 (MOI 10), we measured proinflammatory cytokine/chemokine IL-8, IL-6, CXCL-1, TNF-α, and IL-1β levels in collected supernatants (Fig. 8a-e). UTI89 caused a significant IL-8 (adjusted *p*-value 0.0170) (Fig. 8a), IL-6 (adjusted *p*-value 0.0007) (Fig. 8b), CXCL1 (adjusted *p*-value 0.0004) (Fig. 8c), TNF-α (adjusted *p*-value 0.0340) (Fig. 8d), and IL-1β (adjusted *p*-value 0.0007) response. Although *E. faecalis* 36 showed a significant IL-6 (adjusted *p*-value 0.0092) (Fig. 8b) and TNF-α (adjusted *p*-value 0.0314) (Fig. 8d) release, it caused an IL-8, CXCL-1, and IL-1β response (Fig. 8a, c, e; respectively) that was not statistically significant compared with uninfected control.

**Figure 8.**
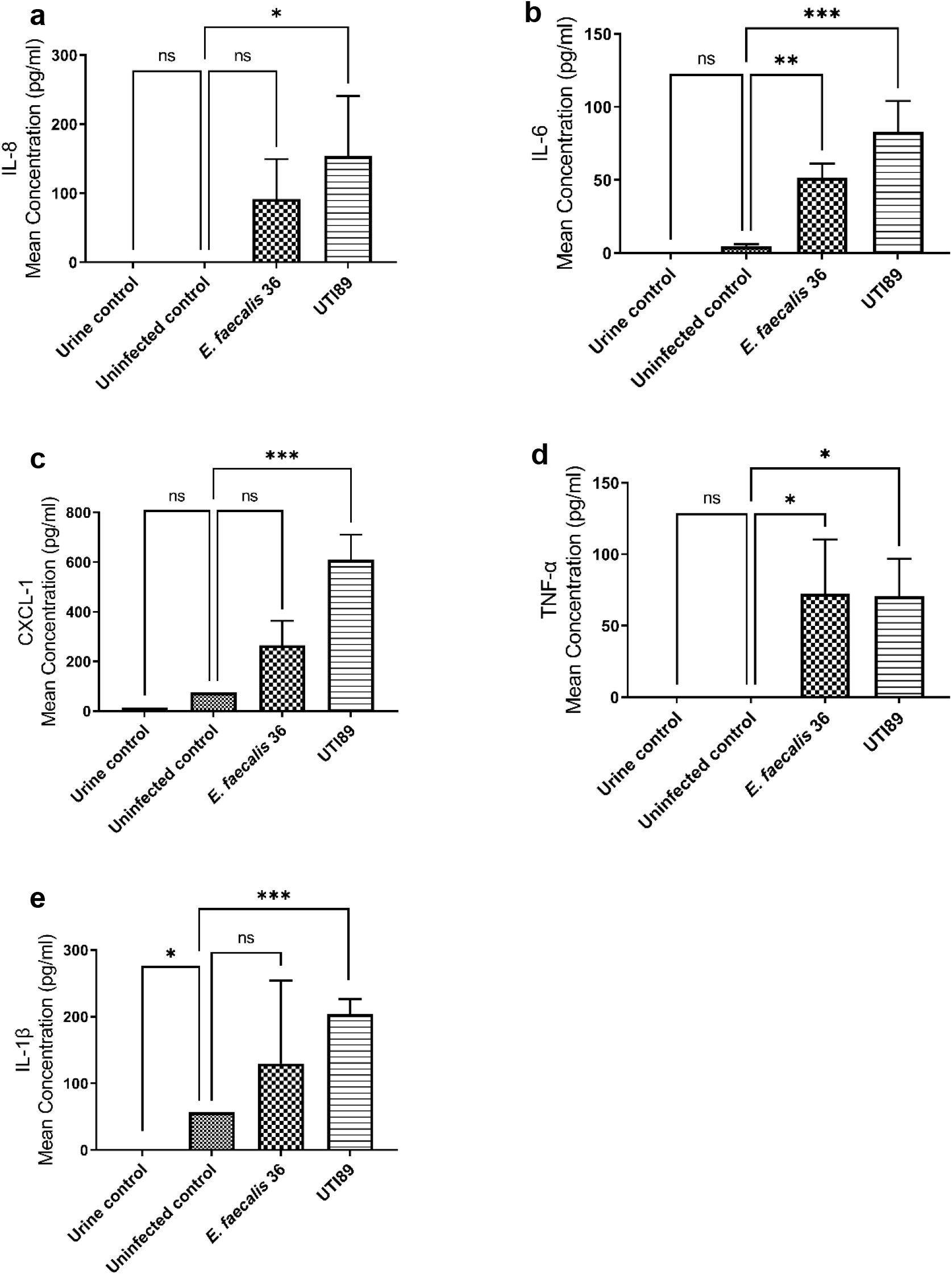
UTI89 and *E. faecalis* 36 trigger pro-inflammatory cytokine/chemokine responses. (a) IL-8; (b) IL-6; (c) CXCL-1; (d) TNF-α, and (e) IL-1β release was measured by ELISA in response to UTI89 and *E. faecalis* 36 strains (MOI 10) at 24h post-infection. Data represent mean ± SD, n=3 biologically independent experiments. *P* values were obtained using ANOVA with Dunnett’s multiple comparisons test.

### Discussion

*In vitro* models of the human urothelium that physiologically resemble human tissue are becoming increasingly available. Such models have improved our understanding of human tissue development and have the potential to complement animal models^32^. Several studies have taken advantage of cells lines and evaluated their capacity to form organoids. Smith *et al*. established a urothelial organoid using 5637 human bladder epithelial carcinoma cell line (HTB-9) under microgravity conditions^33^, while Sharma *et al*. used this cell line to develop a human bladder-chip model^34^. Studies have also described the use of normal human urothelial (NHU) cells to develop a biomimetic urothelial tissue model^35^ and human three-layered bladder assembloids^36^. Here, we present a new advanced barrier-forming, urine-tolerant, 3D urothelial model. This model showed morphological similarities with human bladder urothelium, stratifying to 7 layers (Fig. 1a) with a single basal cell layer at the basal membrane (Fig. 1b), multiple intermediate cell layers and terminally differentiated umbrella-like cells at the apical surface (Fig. 1c). The model also displayed similar CD phenotypes to human bladder tissue^28^, including CD271^+^ basal cells and CD227^+^ umbrella cells (Fig. 2). Although we were able to characterise the cell populations using flow cytometric analysis and specific antibodies, comparing the flow cytometry panel to conventional histochemistry in future will help to distinguish the intermediate cells from other cell layers.

We used immunofluorescence to assess a series of key biomarkers. In doing so, we noticed that the staining of a number of apical markers was discontinuous (Figs. 3-5). We hypothesised that this could be the result of gaps left by recently shed cells, although in some areas the expression of these markers was detectable in focal planes immediately underlying the upper layer, which is very thin. Very little is known, however, about how uniform these markers are in the human urothelium, and future studies are warranted.

Uroplakins are the most prominent markers of urothelial differentiation^37^. We detected all three uroplakins (1A, II, and III) expressed by the terminally differentiated umbrella-like cells in the 3D-UHU model (Fig. 3a-c). Remarkably, our model expressed UPKII, although studies exploring uroplakin expression have stated that in cultured urothelial cells, the glycosylation of pro-UPKII does not occur, thus hampering the formation of the uroplakin heterotetramer leading to impaired AUM assembly^37^ (Fig. 3b). It may be that the stratified and differentiated structure of this 3D model enables the correct environment for this glycosylation to occur as the process is differentiation-dependent^37^. Furthermore, umbrella cells expressed a coating of chondroitin sulfate, a GAG layer marker found in human urothelium (Fig. 3d). Although some components of the GAG layer are difficult to stain by conventional antibodies, further studies into the detailed composition of this layer would be interesting.

Next, we examined the expression of cytokeratins, of which 20 isoforms are known to be expressed by human urothelium^38^. CKs are cytoskeletal polypeptides, and the specific pattern of their expression can be used to determine the cytodifferentiation of the urothelium^31^. Similar to human urothelium, our 3D model exhibited CK8, CK13, and CK20 markers. The 3D-UHU model showed CK8 expression throughout all layers (Fig. 4a) while CK20 was exclusively expressed by terminally differentiated umbrella cells (Fig. 4c). The expression of CK13 has been described by a number of studies^30,31,39^; a late/terminally differentiated transitional phenotype expressing CK13 was detected in the 3D-UHU model (Fig. 4b).

Adherens and tight junctions, together with desmosomes, function as a selective barrier in epithelial cells controlling the paracellular diffusion of water, ions, and various macromoleculars^40^. Cadherins are located on the cell membrane and are important for the function of AJ. We examined the expression of E-cadherin in the 3D-UHU model (Fig. 5a) and observed that it was distributed in the cytoplasm of the apical surface with a faint signal at cell perimeters. In addition, claudins are the main functional barrier-determining component of the TJ^41^ and claudin 3 expression has been shown to associate with differentiation and development of a tight barrier along with ZO-1 protein^42^. The TJs, localised between the umbrella cells, contribute along with uroplakins to urothelial barrier function^12^. However, the intermediate cells, although expressing TJ-associated proteins such as claudins, do not form morphologically identifiable TJ or AJ, which is in contrast to the case in umbrella cells^7^. We demonstrated that ZO-1 expression in 3D-UHU model was localised to the apical surface (Fig. 5b) while claudin 1 and claudin 3 were expressed at apical surface together with diffuse expression throughout the cell layers (Fig. 5 e&f). Although a discontinuous and mainly cytoplasmic expression of the AJ and TJ markers were observed in the 3D-UHU model, examining the markers using sectioned membranes in future may provide an improved staining.

We further examined the TJ formation by TEER assessment. The 3D-UHU model showed a stable increase in TEER, achieving ∼1000 Ω.cm^2^ around day 18-20 (Fig. 5g). Urothelial models generated from two different subcultures (passage 11 & 12) showed a similar trend. Although the 3D-UHU model does not reach the high TEER (usually > 2k Ω.cm^2^) that are reported in models using NHU cell cultures^25,35,43^, unlike the NHU model, 3D-UHU is comprised of several layers, which might affect the TEER values. Further work is required to achieve a somewhat higher TEER value in the presence of full stratification; nevertheless, the moderate barrier function achieved was sufficient to exclude FITC-dextran well and to serve as a baseline for inspecting subsequent loss of barrier function after infection. One study revealed that the barrier integrity of rabbit urothelium mounted in an Ussing chamber was maintained even with a ten-fold drop in TEER as no significant leakage of biotin, fluorescein, or ruthenium was detected across the urothelium^44^. Indeed, when we examined the urothelial barrier function in response to both Gram-positive and Gram-negative uropathogens at MOI of 10 (Fig. 6), we found that CFT073 (a widely studied uropathogenic *E. coli* strain) and *E. faecalis* 36 (a clinical isolate known to be invasive) caused a significant barrier disruption at 24h post-infection.

Finally, we examined some aspects of urothelial innate immunity. First, we identified that HBLAK cells can express TLR-2, TLR-4, and TLR-5 although to varying levels (Fig. 7), all of which are known to be important in urothelial defence in humans. In addition, we wanted to investigate the 3D-UHU innate immune response to infection challenge by two representative uropathogens, UTI89 and *E. faecalis* 36 (Fig. 8). UTI studies in human and mice have reported several cytokine/ chemokine responses to urinary infection and their role in UTI^45,46,47^. A significant IL-8, IL-6, CXCL-1, TNF-α, and IL-1β release was detected in response to UTI89 (Fig. 8a-e). With the *E. faecalis* 36 clinical isolate, infection induced statistically significant IL-6 and TNF-α levels (Fig. 8b & d) similar to that secreted in response to UTI89. This cytokine/chemokine profile suggests that the 3D-UHU model mediates some human-relevant innate immune responses to uropathogenic stimuli, which may make it an appropriate *in vitro* model for understanding innate immune response at the human cell-bacteria interface.

In conclusion, these data taken together suggest that our enhanced 3D-UHU model displays similar structural and physiological features compared with human bladder urothelium and could be used as an *in vitro* model, alongside more traditional animal experiments and human clinical studies, to elucidate UTI host-pathogen interactions.

## Materials and Methods

### Human bladder epithelial progenitor cell culture

HBLAK human bladder progenitor cells (CELLnTEC, Switzerland) were grown according to the CELLnTEC protocol. Briefly, the frozen vial was thawed and transferred to a cell culture vessel containing pre-equilibrated CnT-Prime medium (CnT-PR, CELLnTEC), incubated at 37°C and 5% CO2. HBLAK cells were passaged when reached 70 to 90% confluency and seeded at the recommended density. Cells were propagated to passage 4 then frozen vials were prepared following the CELLnTEC freezing protocol.

### Establishment of 3D-UHU

Human 3D bladder culture was prepared as described previously^48^, with some modifications. Briefly, HBLAK cells were used only within passage 8-12 after thawing. Cells were seeded onto polycarbonate transwell inserts with pore size of 0.4 µm (VWR, United Kingdom) at the recommended seeding density (CELLnTEC, 3D Culture Protocol). An appropriate amount of CnT-PR medium was added to the basolateral chamber of the inserts so that the medium levels were equal. Once cells were confluent (confirmed by staining and sacrificing a parallel well), CnT-PR was replaced with 3D Barrier Medium (CnT-PR-3D, CELLnTEC) and incubated for 15-16h so cells could form intercellular adhesion structures. 3D culture was initiated by replacing apical CnT-PR-3D with filter-sterilized human urine (pooled gender) (BioIVT, United Kingdom) and fresh CnT-PR-3D medium in the basolateral chamber. Urine and CnT-PR-3D medium were changed at regular intervals (twice per week) and 3D cultures were fully stratified and differentiated on days 18-20.

### Characterisation of CD cell surface antigens

To profile 3D-UHU CD cell surface markers, fully differentiated cultures were dissociated using Accutase solution (Merck, United Kingdom). Cells were washed with PBS and filtered (100 µm) to prepare single cell suspension. BD Horizon Fixable Viability Stain 780 solution (1:1000) was added to the cell suspension and incubated at room temperature (RT) for 15 min. Cells were washed and blocked with blocking buffer plus Fc receptor blocking antibody (Clone Fc1.3216, BD Biosciences, United Kingdom). Cells were incubated at RT for 10 min then centrifuged (800 *g*, 3 min). BD Horizon Brilliant Stain Buffer was added to each sample followed by the following antibodies (BD Biosciences): CD9 (BV421, Clone M-L13), CD44 (Alexa Fluor 700, Clone G44-26), CD47 (BV786, Clone B6H12), CD49b (Alexa Fluor 647, Clone AK-7), CD59 (BUV 395, Clone p282), CD63 (PE-Cy7, Clone H5C6), CD95 (BUV737, Clone DX2), CD104 (BV480, Clone 439-9B), CD227 (BV650, Clone HMPV), CD271 (BV711, Clone L128). Cells were incubated at 4 ^°^C in dark for 30 min then washed and resuspended in wash buffer with 1% formaldehyde solution. Data acquisition was performed using Cytek Aurora equipped with 5 lasers.

### Immunostaining of 3D-UHU model and imaging

Prior to immunofluorescence staining, 3D-UHU cultures were fixed with 4% methanol-free formaldehyde (Thermo Fisher Scientific, United Kingdom) overnight at 4 ^°^C. Membranes were cut and placed in a 24-well plate and permeabilised in 0.2% Triton-X100 (Sigma-Aldrich, United Kingdom) in PBS for 20 min at RT. Membranes were washed with PBS then blocked with 5% normal goat serum (Thermo Fisher Scientific) at RT for 1h. 3D cultures were incubated at 4 ^°^C overnight with primary antibodies in 1% bovine serum albumin (BSA)/PBS as follows: rabbit anti-uroplakin-1A (UPK1A) polyclonal antibody (1:100 dilution, PA5-49668, Thermo Fisher Scientific), rabbit anti-uroplakin-II (UPKII) polyclonal antibody (1:100 dilution, NBP2-38904, Novus biologicals); mouse anti-uroplakin-III (UPKIII) monoclonal antibody (1:50 dilution, sc-166808, Santa Cruz); mouse anti-cytokeratin 8 (CK8) monoclonal antibody (1:50 dilution, MA1-06317, Thermo Fisher Scientific); rabbit anti-cytokeratin 13 (CK13) polyclonal antibody (1 µg/ml, PA5-83165, Thermo Fisher Scientific); rabbit anti-cytokeratin 20 (CK20) polyclonal antibody (1:100 dilution, PA5-22125, Thermo Fisher Scientific); rabbit anti-E-cadherin polyclonal antibody (1:200 dilution, PA5-32178, Thermo Fisher Scientific); rabbit anti-zonula occludens-1 (ZO-1) polyclonal antibody (2.5 µg/ml, 40-2200, Thermo Fisher Scientific); mouse anti-claudin 1 monoclonal antibody (2 µg/ml, 37-4900, Thermo Fisher Scientific); rabbit anti-claudin 3 polyclonal antibody (1:100 dilution, PA5-16867, Thermo Fisher Scientific); mouse anti-chondroitin sulfate antibody (1:100 dilution, ab11570, Abcam). HBLAK cells grown on membranes were incubated with the following primary antibodies: rabbit anti-TLR-2 polyclonal antibody (10 µg/ml, PA5-20020, Thermo Fisher Scientific), rabbit anti-TLR-4 polyclonal antibody (5 µg/ml, PA5-23124, Thermo Fisher Scientific), and rabbit anti-TLR-5 polyclonal antibody (10 µg/ml, 36-3900, Thermo Fisher Scientific).

Post incubation, membranes were washed with 1% BSA/PBS and incubated with 1:500 dilution of secondary antibodies (goat anti-mouse or goat anti-rabbit) conjugated with Alexa Fluor-488 (Thermo Fisher Scientific) at RT for 1.5 h. Membranes were washed then stained with Alexa Fluor-633 phalloidin (1:500 dilution, Thermo Fisher Scientific) to label filamentous actin followed by DAPI nucleic acid stain (DAPI dihydrochloride, 300 nM in PBS, Invitrogen, United Kingdom). Filter membranes were washed then mounted with ProLong Glass antifade mountant (Invitrogen) and imaged on a Leica SP8 microscope.

To analyse urothelial cell shedding, apical supernatants from a 12-well plate was collected and centrifuged at 300 *g* for 5 min. Pellets were re-suspended in 100 µl of PBS and cytocentrifuged onto a glass slide using a Shandon Cytospin 2 at 800 rpm for 5 min. Cells were fixed with 4% methanol-free formaldehyde for 15 min at RT then washed 3x in Hanks′ Balanced Salt solution (HBSS) (Thermo Fisher Scientific). Cells were stained with 5 µg/ml of Wheat Germ Agglutinin (WGA), Alexa Fluor-488 conjugate (Thermo Fisher Scientific) for 1h at RT. Cells were washed with HBSS and stained with 2 µg/ml Hoechst (33342, Thermo Fisher Scientific) for 15min at RT, and mounted as described above.

### Transepithelial electrical resistance (TEER) and paracellular permeability measurements

The TEER in 3D-UHU model was measured using EVOM3 with STX2-Plus electrode (World Precision Instruments, United Kingdom) according to the manufacturer’s instruction. Briefly, 1000Ω test resistor was used to set up the equipment then blank handling mode was selected to allow subtraction of the blank control from the current resistance measurement of 3D cultures. Each measurement was recorded and stored on the device for further analysis.

Barrier integrity of 3D cultures were assessed using Fluorescein Isothiocyanate (FITC)-Dextran (MW 4,000, Sigma-Aldrich, United Kingdom). FITC-dextran solution (1mg/ml) was prepared in urine and added to the apical chamber. Aliquots were collected in 0, 5, and 24h post-infection (multiplicity of infection; MOI 10) and fluorescent was measured by a Tecan microplate reader (Spark, Switzerland) at an excitation of 485 nm and emission of 538 nm. The FITC-conjugated dextran concentration was presented as relative fluorescence unit (RFU).

### Bacterial strains

To examine the barrier disruption and innate immune response to bacterial co-culture, several uropathogenic strains was employed. Uropathogenic *Escherichia coli* (UPEC) strains UTI89^49^, CFT073^50^ (ATCC 700928), and *E. coli* 83972 (bei Resources) associated with asymptomatic bacteriuria (ABU) were grown in static Luria-Bertani (LB, Sigma Aldrich) broth at 37°C for 48h to induce expression of type 1 pili. Clinical isolates of *Enterococcus faecalis* (*E. faecalis* 36 [EF36], previously isolated and reported from a patient with chronic UTI^51^, and *E. faecalis* 77^52^, previously isolated and reported from an asymptomatic healthy male) and *Streptococcus agalactiae*, previously isolated and reported^52^ (Group B Streptococcus), were grown overnight in Tryptone Soya Broth (TSB, Thermo Fisher Scientific) at 37°C.

### Co-culture and cytokine measurements

The 3D-UHU were co-cultured with uropathogenic isolates at MOI 10. To measure the expression of pro-inflammatory cytokines/chemokines, the apical supernatants were collected at 24h post infection. Human Interleukin-8 (IL-8), IL-6, IL-1β, and TNF-α was measured by enzyme-linked immunosorbent assay (ELISA) per manufacture’s instruction (Biolegend, ELISA MAX Deluxe Set) and CXCL-1/GRO-α was measured using a R&D systems DueSet ELISA kit.

### Statistical analysis

Data were analysed using GraphPad Prism 9. Three independent repeats in duplicates were performed for statistical analysis. Differences in FITC-dextran measurements and cytokine/chemokine expression between experimental samples and controls were analysed using 2way-ANOVA. Dunnett’s multiple comparisons test was used to compare each variable with the untreated control.

## Acknowledgements

We would like to thank the members of our laboratory for helpful discussions, and a philanthropic donation (UCL 540059) for funding this work. UTI89 was a kind gift from Prof. Scott Hultgren’s lab.

